# Structural and molecular dynamics of *Mycobacterium tuberculosis* malic enzyme, a potential anti-TB drug target

**DOI:** 10.1101/2020.07.07.192161

**Authors:** Kalistyn H. Burley, Bonnie J. Cuthbert, Piyali Basu, Jane Newcombe, Ervin M. Irimpan, Robert Quechol, Ilona P. Foik, David L. Mobley, Dany J.V. Beste, Celia W. Goulding

**Author notes:** Contributed Equally (Kalistyn H. Burley, Bonnie J. Cuthbert).

## Abstract

Tuberculosis (TB) is the most lethal bacterial infectious disease worldwide. It is notoriously difficult to treat, requiring a cocktail of antibiotics administered over many months. The dense, waxy outer membrane of the TB-causing agent, *Mycobacterium tuberculosis* (Mtb), acts as a formidable barrier against uptake of antibiotics. Subsequently, enzymes involved in maintaining the integrity of the Mtb cell wall are promising drug targets. Recently, we demonstrated that Mtb lacking malic enzyme (MEZ) has altered cell wall lipid composition and attenuated uptake by macrophages. These results suggest that MEZ provides the required reducing power for lipid biosynthesis. Here, we present the X-ray crystal structure of MEZ to 3.6 Å resolution and compare it with known structures of prokaryotic and eukaryotic malic enzymes. We use biochemical assays to determine its oligomeric state and to evaluate the effects of pH and allosteric regulators on its kinetics and thermal stability. To assess the interactions between MEZ and its substrate malate and cofactors, Mn^2+^ and NAD(P)^+^, we ran a series of molecular dynamics (MD) simulations. First, the MD analysis corroborates our empirical observations that MEZ is unusually disordered, which persists even with the addition of substrate and cofactors. Second, the MD simulations reveal that MEZ subunits alternate between open and closed states and that MEZ can stably bind its NAD(P)^+^ cofactor in multiple conformations, including an inactive, compact NAD^+^ form. Together the structure of MEZ and insights from its dynamics can be harnessed to inform the design of MEZ inhibitors that target Mtb.

---

Tuberculosis (TB) is the leading cause of infectious death globally with 1.5 million TB-related deaths and 1.1 million new infections in 2018 alone [1]. This global pandemic is fueled by the emergence of drug-resistant strains of the causative agent *Mycobacterium tuberculosis* (Mtb) as well as a catastrophic synergy with HIV. TB treatment involves a cocktail of antibiotics which need to be taken for a minimum of 6 months [2]. Non-compliance is common and has contributed to the emergence of multi- and extensively-drug resistant Mtb strains. New, less toxic anti-TB therapeutics with shorter treatment regimens are urgently needed.

In a recent study we demonstrated that the enzymes at the anaplerotic node of central carbon metabolism are essential for Mtb intracellular survival in macrophages and therefore are potential therapeutic targets [3]. The anaplerotic node consists of enzymes that connect the pathways of glycolysis, gluconeogenesis and the tricarboxylic acid (TCA) cycle. These proteins regulate the flux distribution between anabolism, catabolism and energy supply to the cell, while replenishing the intermediates of the TCA cycle. One of the enzymes in this node is malic enzyme (ME), which can reversibly convert malate and NAD(P)^+^ to pyruvate, CO_2_ and NAD(P)H. Our previous study suggests that Mtb ME (MEZ) has a role in lipid biosynthesis (forward direction as a L-malate decarboxylase) and a minor back-up role in CO_2_-dependent anaplerosis (reverse reductive pyruvate carboxylation reaction), **Figure S1** [3]. Although we showed that MEZ is not required for growth on gluconeogenic or glycolytic substrates, the MtbΔ*mez* deletion strain had impaired macrophage invasion [3]. Additionally, the MtbΔ*mez* colonies had a glossy and viscous morphology, which has been observed previously for Mtb mutants that lack cell-wall lipid transporters and a trehalose dimycolate esterase [4-7], suggesting that the cell-wall lipid composition had been altered. Indeed, the MtbΔ*mez* strain had a buildup of apolar free fatty acids and mycolic acids as compared to wildtype Mtb, which resulted in an accompanying decrease in the levels of cell-wall bound mycolates in total lipid extractions [3]. These results suggest that a major role of Mtb MEZ is as a L-malate decarboxylase, to produce NAD(P)H for lipid biosynthesis.

It has been shown that *mez* is part of the *phoPR* regulon, a two-component system that regulates genes essential for the biosynthesis of complex lipids, which supports the role of MEZ in lipid metabolism [8]. Specifically, the MtbΔ*phoP* mutant, similar to MtbΔ*mez*, results in a structurally altered cell envelope composition providing further evidence that MEZ supports NAD(P)H production for the synthesis of complex lipids. The connection between MEs and lipid biosynthesis has been observed in other species [9-11]. Studies of eukaryotic oleaginous microorganisms, with an emphasis on algae and yeast, demonstrate that MEs produce NAD(P)H for lipid biosynthesis [10] including fatty-acid rich storage compounds such as triacylglycerol (TAG). In *Streptomyces coelicolor*, it has been shown that deletion of the NAD^+^- and NADP^+^-dependent ME expressing genes results in the decreased production of TAG [9]. Finally, the overexpression of ME in *Rhodococcus jostii* during growth on glucose promoted an increased NADP^+^-dependent ME activity resulting in a 2-fold increase in fatty acid biosynthesis [11]. Together these studies support that MEZ plays a major role in the production of reducing agent required for Mtb lipid biosynthesis and virulence, and therefore it is compelling to consider MEZ as a potential candidate for structure-based anti-TB drug design.

The biological importance of MEs is reflected in its ubiquity amongst many species, including eukaryotes, prokaryotes and archaea. It follows that there is a large, diverse group of X-ray crystal structures of NADP^+^- and NAD^+^-dependent MEs in the Protein Data Bank (PDB, **Table S1 & Figure S2A**) ranging from high eukaryotes to prokaryotes, all of which contain a NAD(P)H-binding Rossman-fold domain [12-14]. Characterized MEs typically adopt dimeric or tetrameric assemblies with subunits ranging from 40-50 kDa for smaller prokaryotic MEs to ∼60 kDa for many larger subunit MEs found in both eukaryotes and prokaryotes (**Figure 1A**). To-date, there have been three distinct structural classes of MEs reported: small subunit, large subunit and chimeric (the subset of chimeric MEs will not be further discussed). The structures of eukaryotic large subunit MEs, consist of a tetrameric assembly comprised of a dimer of dimers (**Figure 1B**). Each monomer has four distinct domains (A-D). Within higher eukaryotes, the presence of a C-terminal extension in Domain D facilitates the tetramerization of two dimers [14]. By contrast, in plant MEs, dimer tetramerization is facilitated by an N-terminal extension in Domain A [12]. There is one large subunit prokaryotic ME in the PDB from *Escherichia coli* (PDB: 6AGS) with a dimeric, non-tetrameric assembly. The other distinct class consists of prokaryotic small subunit MEs, which are dimeric and have a minimal Domain A and no Domain D. For these small subunit MEs assembly of the dimer is required for full formation of the active site, wherein each monomer subunit contributes a Lys and a Tyr to the active site that are required for catalysis [13]. By comparison, large subunit ME proteins have fully formed active sites, independent of dimerization. Alignment of representative prokaryote and eukaryote large subunit ME sequences highlight the presence of an extended N- and/or C-terminal extension among the larger subunit MEs that is unique to eukaryotes and is required for tetramer formation (**Figure 1B)**. In all structural classes of MEs reported, Domains B and C have conserved motifs and a high degree of structural similarity. A large body of biochemical data exists for MEs, particularly for higher eukaryotic and plant MEs, where ME mechanism of action, catalytic residues and activators and inhibitors have been well studied [12, 15-18]. This has been particularly well-reviewed for higher eukaryotes by Tong *et al*. [14]. In contrast, there is much less known about the biochemistry of large subunit prokaryotic MEs, apart from *E. coli* MEs [19, 20].

**Figure 1.**
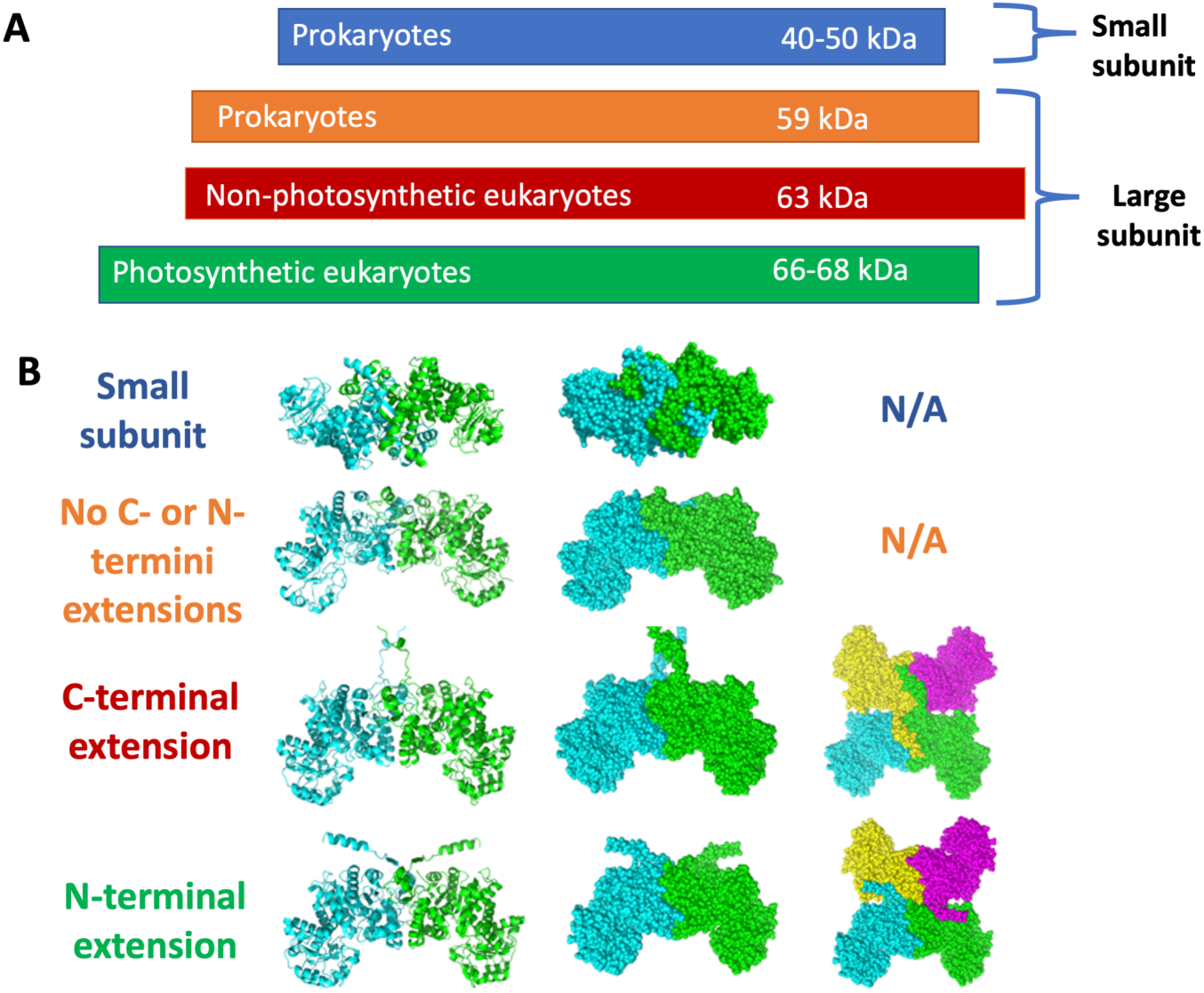
Malic Enzyme Structural Classes and Oligomeric States. **A**. MEs can be classified as small subunit (blue) and large subunit (orange, red, green). Small subunit MEs are typically prokaryotic, while large subunit MEs have been found in both prokaryotes and eukaryotes. Among large subunit MEs, the presence and/or absence of N- and C-terminal tails broadly corresponds with its source species. **B**. Small subunit MEs form dimers (PDB: 1WW8). Among large subunit MEs, slight variations are observed in the lengths of the N- and C-terminal tails. Tetrameric models of human mitochondrial ME (C-terminal extension, PDB: 3WJA), and maize ME (N-terminal extension, PDB: 5OU5) reveal how the extended termini interact with subunits of neighboring dimers to stabilize the formation of the tetramer. *E. coli* ME (PDB: 6AGS), represented in the second row, has neither an N-nor C-terminal extension and exists as a dimer.

The X-ray crystal structure of MEZ is of paramount importance to understanding the enzyme’s biochemistry and assessing its suitability as an anti-TB drug target. In this study, we solved the structure of MEZ in its apo form and determined that MEZ, even though it is a member of the large subunit ME family, is dimeric. Previously, we demonstrated that MEZ utilizes NAD^+^ or NADP^+^ as its cofactor [3]. To shed light on the divalent metal, substrate and cofactor binding modes, we carried out molecular dynamics (MD) simulations on the docked MEZ dimer structure. These simulations highlighted several unique factors about MEZ; (1) the requirement of divalent metal for malate binding; (2) the overall inherent structural flexibility of MEZ compared to other ME structures; and (3) the most stable MEZ dimer appears to form a mixed open: closed conformation in the presence of Mn^2+^, malate and NAD(P)^+^. Finally, we highlight sequence, structural and cofactor binding mode differences between MEZ and human ME, which can be harnessed as a template for MEZ structure-based inhibitor design. Altogether, this work contributes significantly to the biochemical and structural understanding of MEZ and establishes the potential of this enzyme as an anti-TB drug target.

## Results

### Structure of Mtb Malic Enzyme (MEZ)

The asymmetric unit contains four subunits or two dimers of MEZ. Dimer 1 is comprised of subunits A and B, while dimer 2 is formed by subunits C and D. Dimer 1 and dimer 2 are very similar with a root mean square deviation (rmsd) of 0.88 Å over all αC atoms. Similarly, the subunits align closely, where subunit A is most like subunit D with an rmsd of 0.72 Å and subunit B is most like subunit C with an rmsd of 1.05 Å. In the dimers, subunit A aligns to subunit B with an rmsd of 1.37 Å, and subunit C aligns to subunit D with an rmsd of 1.12 Å.

Like other MEs [14, 17, 21], MEZ is composed of four domains (**Figures 2A & B**). Domain A (residues 1-113) is composed of five α-helices (α1-α5, **Table S2**). Domain B (residues 114-262, and 455-525) has an internal β-sheet composed of 5 parallel β-strands (β1/β5/β2/β6/β7). The rest of domain B is helical: α6-α12 and α20-α23. Domain C is formed by residues 263-454, and bears the canonical Rossmann fold of ME proteins. The Rossmann-fold has a 6-stranded parallel β-sheet (β9/β8/β10/β11/β12/β14). Preceding the final β-strand of the β-sheet is a conserved β-hairpin composed of strands β13 and β14. The β-sheet is enclosed by seven α-helices: α13-α19. Lastly, the tail of the C-terminus (residues 526-545) composes domain D, which is largely unstructured except for a final α-helix: α24 or α25.

**Figure 2.**
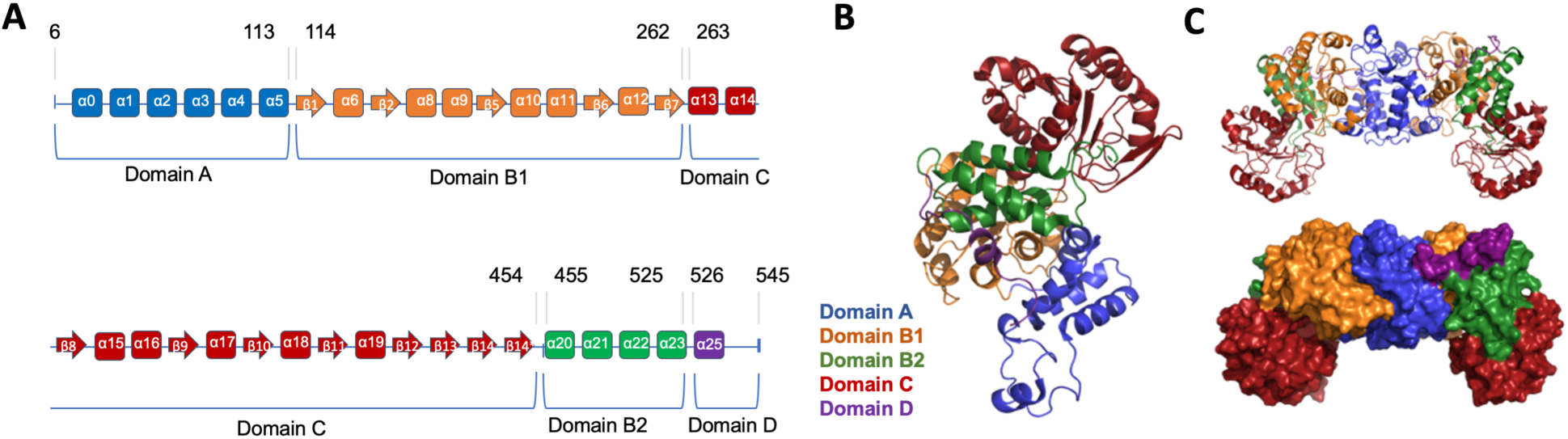
Structural Domains of MEZ. **A**. The structural domains of MEZ with corresponding residue numbers at the start and end of each domain. β-strands and α-helices are represented with arrows and rectangles, respectively. **B**. Structural cartoon of the MEZ monomer with the domains colored as in **A. C**. Structural surface representation of dimeric MEZ with domains colored as in **A**.

The four subunits of MEZ are very similar. However, there are some noteworthy differences, **Table S2**. There are variations in secondary structure between individual subunits; for example, β-strands β3 and β4 are absent in subunit A, but present in subunits B, C and D. Additionally, α-helix α7 is present in only subunit B, α17 is present in every chain except B, α21 is present only in subunit A, α24 is only present in subunit B, and α25 is only present in subunits A and C. These differences between subunits underlie the overall disorder that is observed in the MEZ structure. A rather unexpected difference is that within subunit B, the proposed Rossmann-fold NAD(P)H binding site is not modeled due to an absence of observable electron density and is likely unstructured, potentially indicating increased mobility in this region compared to other MEs. Additionally, there is some variability in modeled α-helices between subunits. Subunit C has an additional helix (residues 11-15) before α1, which aligns to an α-helix in human ME (PDB: 1QR6) [18]. All subunits show a substantial kink in α5, where, in subunit D, the α-helix is interrupted between residues 96-102; similar to subunit D, a break is also observed in the structure of human ME (residues 111-113).

More striking is the difference in the β-sheet strands between subunits. ME proteins have a conserved β-hairpin that is not present in subunit A, but is observed in subunit C (β3/β4) and is partially formed in subunits B and D. Additionally, in the Rossmann fold β-sheet, strand β6 is interrupted in subunit C due to a complete loss of density. Lastly, while the Rossmann fold of subunit A shows the canonical 6-stranded β-sheet with residues 451 and 452 forming an abridged β-strand (β14*) following β-hairpin strand β14, in the other subunits these residues form a continuous β-strand (β14) with varying likelihoods of participating in the Rossmann fold. Notably, in chains B and C, β14 is shifted away from the Rossmann fold β-sheet, however, in subunit B, it is unclear whether either β13 or β14 participate in the β-sheet. While in subunit D, β14 aligns well to β14* positioning and is likely to participate in the Rossmann fold β-sheet. This suggests that, in some MEZ conformations, the Rossmann-fold β-sheet stabilizes the β-hairpin (β13/β14) and reduces its mobility.

Interestingly, in β9 there is a defined kink that is not observed in other MEs due to MEZ Pro328, **Figures S2B & S3**. When MEZ is superimposed onto human ME [18] (PDB: 1QR6) or ‘aster yellows witches’-broom’ *Candidatus* Phytoplasma (AYWB) ME [13] (PDB: 5CEE) structures, measuring the angle between the α-carbon of P328 to the α-carbons of human ME I341 and M343, and from α-carbon of P328 to the α-carbons of AYMB ME V341 and L213 gives the resulting angle of 6° and 17°, respectively. This kink results in Arg331 being oriented such that the NAD(P)^+^ cofactor is stabilized within the active site (**see Figure 6**), further discussed below.

The MEZ structure shows an ‘open’ conformation, which is unsurprising considering the divalent metal, NAD(P)H and substrate-binding sites are unoccupied except for a glycerol molecule from the cryoprotectant in subunit A. Notably, compared to other ME structures in the PDB, MEZ appears to be the least ordered as (1) many secondary structure elements are random coils or have no observable electron density in the MEZ structure compared to others and (2) MEZ appears to be relatively unstable as considerable efforts were made to improve the diffraction quality of MEZ crystals to no avail.

### The oligomeric state of MEZ

Most of the reported large subunit ME proteins are tetrameric [12, 14]. As MEZ falls into the large subunit category, it was surprising that the MEZ crystal structure is dimeric. The MEZ X-ray crystal structure has four subunits in the asymmetric unit, where the two dimers have a similar assembly to the large subunit ME dimers (**Figure 2C**). Whilst the two dimers do weakly interact to form a tetramer, the MEZ tetrameric state does not resemble any previously observed ME tetramer assemblies, **Figure S2C**. Furthermore, the MEZ sequence does not have an N-or C-extension that facilitates tetramer formation in plant and higher eukaryote MEs, respectively (**Figure S2B**).

The dimer interface, similar to those seen in other large subunit ME proteins, is facilitated predominately by hydrophobic interactions and some H-bonds between Domain A from each monomer. Additionally, Domain A from one monomer also interacts with a couple of regions in Domain B from the other monomer. The surface areas buried in Dimer 1 and Dimer 2 are 2431 and 2428 Å^2^, respectively, where the monomers have a surface area of 24381-27347 Å^2^. At the dimer/dimer interface of the MEZ structure there is very little surface area buried, 396 and 379 Å^2^ for Subunits A & D and B & C, respectively, suggesting that the tetramer in the asymmetric unit is a consequence of crystal packing rather than MEZ forming a stable tetramer in solution. Strikingly, the only other prokaryotic large subunit ME structure (PDB: 6AGS, *E. coli* ME) is also dimeric. The *E. coli* ME dimer superimposes well with the MEZ dimer (rmsd 2.20 Å over all atoms) and has a similar dimer interface. Notably, in the *E. coli* ME dimer, the first and last β-strand from alternate monomers form two 2-stranded β-sheets, in contrast to the MEZ structure, where the N- and C-termini of all monomers are random coils (**Figure 2**). To confirm the oligomeric state of MEZ in solution, we carried out size exclusion chromatography (SEC) and the results suggest that MEZ is predominately dimeric in solution at pH 7.0 (**Figure 3A**). Thus, both structurally characterized large subunit bacterial MEs appear to be dimeric, although biochemical data of *E. coli* ME suggests it assembles as a tetramer [20].

**Figure 3.**
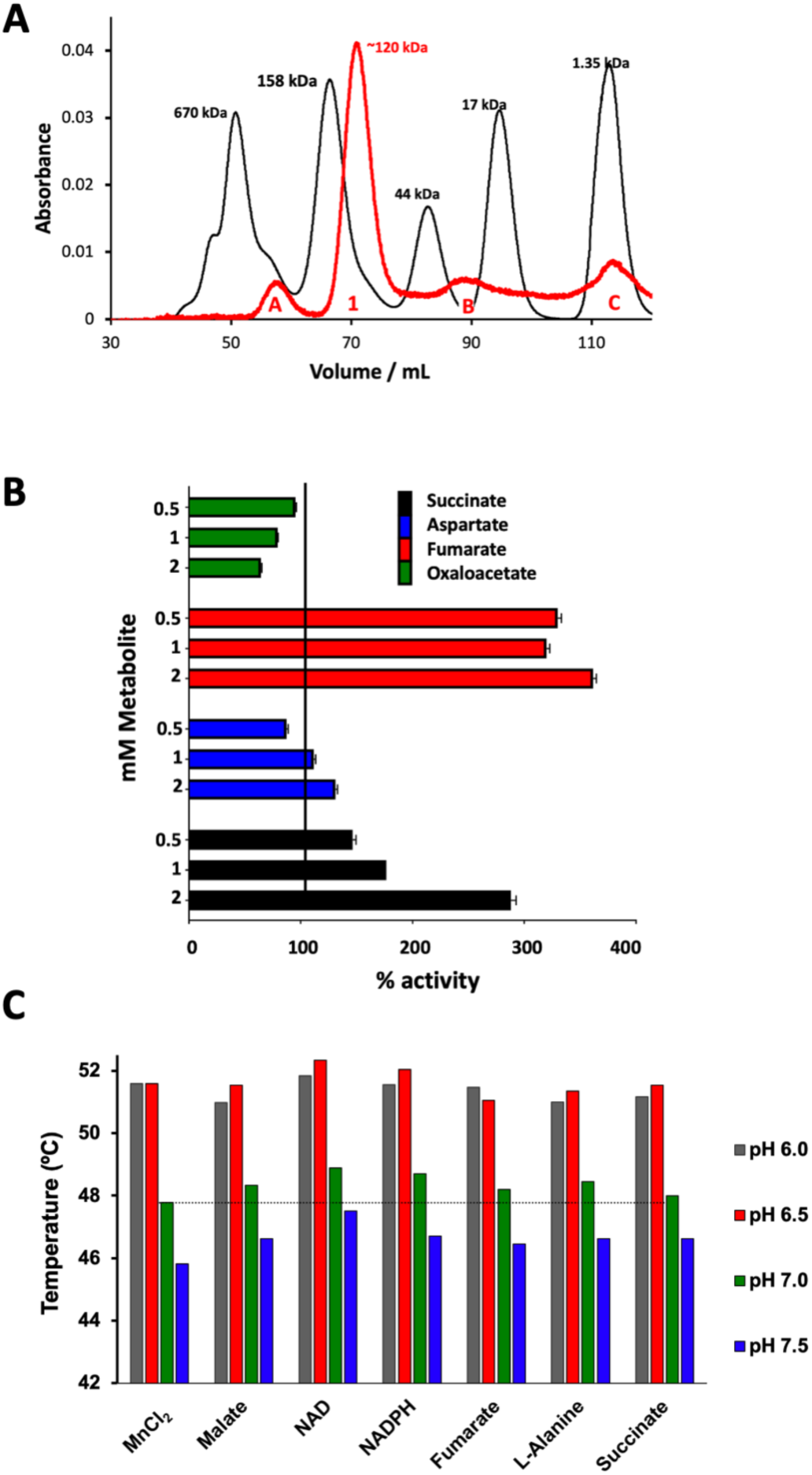
Biophysical and Biochemical Analyses of MEZ. **A**. Gel filtration chromatogram of MEZ to identify its oligomeric state. His-tagged MEZ was run on a S200 HiLoad 16/600 column at 0.5mL/min, and compared with protein standards. The predominant MEZ peak (Peak 1) is the dimeric species (expected MW is 124 kDa), which runs at an approximate size of 120 kDa. SDS-PAGE analysis of peak B reveals a contaminant protein, and peak C is probably imidazole. Peak A is likely higher order MEZ aggregates. **B**. Regulatory properties of recombinant MEZ. MEZ forward activity using NAD^+^ as a cofactor was measured at pH 7.4 in the absence or presence of each effector whilst the malate concentration was maintained at 4 mM. Values are expressed as percentages of the activity measured in the control assay containing no effector. **C**. Thermal stability (*T*_*m*_) of MEZ measured at various pHs with different small molecules (2mM) using DSF. All conditions included 2mM MnCl_2_. Experiments were performed at least in triplicate.

### The kinetics and allosteric regulation of MEZ

We previously showed that, although MEZ was able to catalyze the reverse reductive pyruvate carboxylation, this enzyme clearly has a strong preference for the forward gluconeogenic direction [3]. To inform the structural studies, we performed a more detailed kinetic characterization of purified MEZ at two different pHs, **Table 2**. At pH 7.4 the initial rates of MEZ measured in various concentrations of malate demonstrated sigmoidal kinetics irrespective of the co-factor used, which implies positive cooperative malate binding and has been described for other ME’s [22, 23] Although the Hill coefficients were similar when either cofactor was used, the *K*_0.5_ for malate (NAD^+^) was approximately 50% lower than the *K*_0.5_ for malate (NADP^+^). The cooperative effect of malate binding was enhanced at pH 6.6 in the presence of NAD^+^ but was abolished with NADP^+^ as the cofactor. Furthermore, due to enhanced malate cooperativity at low pH, we also observe an increase in catalytic efficiency of MEZ with NAD^+^ at pH 6.6. The results also suggest that binding of NAD^+^ occurs in a positive cooperative fashion whereas the binding of NADP^+^ follows Michaelis-Menten kinetics. Cooperativity would enable MEZ to produce step-like rate responses to changes in malate and NAD^+^ concentration, and thereby respond rapidly to changing cellular conditions. We confirmed that MEZ preferred NAD^+^ over NADP^+^, with a *K*_*0*.*5*_ value for NAD^+^ nearly 3 times lower than the *K*_*m*_ for NADP^+^ at pH 7.4 (**Table 2**).

**Table 1.** Data collection and refinement statistics for MEZ crystal structure.

|  | MEZ |
| --- | --- |
| **Data collection** |  |
| Space group | P 2<sub>1</sub> |
| Cell dimensions |  |
| *a*, *b*, *c* (Å) | 94.6, 144.1, 118.1 |
| α, β, γ (°) | 90, 109.5, 90 |
| Resolution (Å) | 89.2-3.6 (3.7-3.6)<sup>a</sup> |
| *R*<sub>merge</sub><sup>b</sup> | 0.178 (0.917) |
| *I* / σ*I* | 2.9 (0.87) |
| Completeness (%) | 97.62 (94.86) |
| Redundancy | 2.0 (1.9) |
| **Refinement** |  |
| Resolution (Å) | 89.2-3.6 (3.7-3.6) |
| Total reflections | 66726 (6289) |
| Unique reflections | 33948 (3301) |
| *R*<sub>work</sub> / *R*<sub>free</sub><sup>c</sup> | 32.1 / 35.8 |
| Ramachandran favored (%) | 90.0 |
| Ramachandran outliers (%) | 0.25 |
| No. atoms |  |
| Protein | 13575 |
| Ligands | 72 |
| Water | 145 |
| *B*-factors (Å<sup>2</sup>) |  |
| Protein | 86.31 |
| Ligands | 86.78 |
| Water | 54.01 |
| R.m.s. deviations |  |
| Bond lengths (Å) | 0.01 |
| Bond angles (°) | 1.06 |
| **PDB-ID** | **6URF** |

<sup>a</sup> Values within parentheses refer to the highest resolution shell.
<sup>b</sup> $R_{\text{merge}} = \sum \sum |I_{\text{hkl}} - I_{\text{hkl}}(j)| / \sum I_{\text{hkl}}$ , where $I_{\text{hkl}}(j)$ is observed intensity and $I_{\text{hkl}}$ is the final average value of intensity. $R_{\text{pim}} = \sum \{1/[N_{\text{hkl}}-1]\}^{1/2} \times \sum |I_{\text{hkl}}(j) - I_{\text{hkl}}| / \sum \sum I_{\text{hkl}}(j)$ . $R_{\text{meas}} = \sum \{N_{\text{hkl}}/[N_{\text{hkl}}-1]\}^{1/2} \times \sum |I_{\text{hkl}}(j) - I_{\text{hkl}}| / \sum \sum I_{\text{hkl}}(j)$ . $\text{CC} = \sum (x-(x))(y-(y)) / [\sum (x-(x))^2 \sum (y-(y))^2]^{1/2}$ .
<sup>c</sup> $R_{\text{work}} = \sum |F_{\text{obs}}| - |F_{\text{calc}}| / \sum |F_{\text{obs}}|$ and $R_{\text{free}} = \sum |F_{\text{obs}}| - |F_{\text{calc}}| / \sum |F_{\text{obs}}|$ , where all reflections belong to a test set of 5% data randomly selected in Phenix.

**Table 2.** Kinetic constants for MEZ.

| Substrate/cofactor (pH) | $K_m$ or $K_{0.5}$<br>(mM) | $V_{max}$<br>(mmol min<sup>-1</sup> µg<sup>-1</sup>) | $V_{max}/K_m$ or $K_{0.5}$<br>(min<sup>-1</sup>µg<sup>-1</sup>) | Hill<br>coefficient |
| --- | --- | --- | --- | --- |
| **Gluconeogenic reaction**<br>**(L-malate decarboxylation)** |  |  |  |  |
| <sup>a</sup>L-malate with constant NAD<sup>+</sup> (pH 6.6) | 3.92±0.17<sup>b</sup> | 63.15±0.18 | 16.11 | 1.6-2.2 |
| <sup>a</sup>L-malate with constant NADP<sup>+</sup> (pH 6.6) | 16.6±1.34 | 43.97±1.05 | 2.65 |  |
| <sup>a</sup>L-malate with constant NAD<sup>+</sup> (pH 7.4) | 18.86±0.52<sup>b</sup> | 127.3±1.75 | 6.75 | 2.1-2.7 |
| <sup>a</sup>L-malate with constant NADP<sup>+</sup> (pH 7.4) | 36.85±2.22<sup>b</sup> | 48.27±2.29 | 1.31 | 2.2-2.6 |
| <sup>c</sup>Constant L-malate with NAD<sup>+</sup> (pH 6.6) | 0.49±0.35<sup>b</sup> | 16.55±0.44 | 33.77 | 1.4-1.6 |
| <sup>c</sup>Constant L-malate with NADP<sup>+</sup> (pH 6.6) | 0.76±0.13 | 58.65±4.8 | 77.17 |  |
| <sup>c</sup>Constant L-malate with NAD<sup>+</sup> (pH 7.4) | 0.50±0.03<sup>b</sup> | 16.83±0.51 | 33.66 | 1.4-1.6 |
| <sup>c</sup>Constant L-malate with NADP<sup>+</sup> (pH 7.4) | 1.43±0.02 | 21.14±0.22 | 14.78 |  |

Kinetic values are given as averages ± SEM.
<sup>a</sup>L-malate (0.25 - 100 mM) and NAD(P)<sup>+</sup> is 1.6 mM
<sup>b</sup>Kinetic for these reactions are sigmoidal and the reported values are $K_{0.5}$ values
<sup>c</sup>NAD(P)<sup>+</sup> (0.05 - 1.6 mM) and [L-malate] is 100 mM

Previous studies have shown that several metabolic intermediates can act as allosteric regulators of MEs [15,19,24-27]. When NAD^+^ is the MEZ cofactor, the addition of fumarate and succinate activated the forward malate decarboxylation reaction to 361% ad 290% of the unmodified activity, respectively, and aspartate induced a smaller increase in activity, **Figure 3B**. Like *E. coli* ME, oxaloacetate inhibited MEZ activity to 61% [19], **Figure 3B**. Finally, pyruvate, glutamate and acetate did not significantly modify the MEZ activity even at high concentrations (data not shown).

To evaluate the effects of pH and potential allosteric regulators on the oligomeric assembly of MEZ, we performed SEC analysis at pH 6.0 and pH 7.4 after incubation of MEZ with 100-fold molar excess of fumarate, succinate and alanine. In all conditions studied, MEZ remained predominately dimeric (data not shown) as observed for MEZ alone (**Figure 3A**).

We further examined the impact of ligand binding and pH on the thermal stability of MEZ. Using differential scanning fluorimetry (DSF), we determined the melting temperature (*T*_*m*_) of MEZ in the presence of 10 mM MnCl_2_ with 200-fold molar excess of NAD(P)^+^, malate, or potential allosteric activators at various pHs (6.0, 6.5, 7.0 and 7.5). The addition of cofactors or substrate had minimal impact on the *T*_*m*_ of MEZ, with the greatest shift (∼1.7°C) occurring at pH 7.5 between apo MEZ and MEZ in the presence of NAD^+^ (**Figure 3C**). While incubation with potential ME allosteric activators had little effect, there is a clear trend between MEZ thermal stability and pH. At low pH (6.0 & 6.5), the *T*_*m*_s are comparable, ranging from 51.0-52.5°C; at higher pH (7.0 & 7.5), the measured *T*_*m*_s are lower, ranging from 45.5-49°C. MEZ is stabilized in a more acidic environment and this increased stability coincides with elevated activity in the kinetic experiments (**Table 2**). Thus, MEZ appears to be partially regulated by pH: destabilized and less active at higher pH, and more stable and active at lower pH. Similarly, the activity of photosynthetic MEs are regulated by pH [12].

### The divalent metal and active site

All MEs studied thus far, require binding of either Mn^2+^ or Mg^2+^ in the active site to reinforce the structural integrity of the enzyme and to aid in substrate, intermediate and product binding during catalysis [28, 29]. The divalent metal binding site is highly conserved among both prokaryotic and eukaryotic MEs and comprises two Asp residues and one Glu residue: this conservation is also observed in MEZ and consists of Glu240, Asp241, and Asp264 (**Figure S2B**). Additional critical catalytic residues in other MEs studied are Lys169 and Tyr96 (numbering for MEZ) [14]. In eukaryotes, an Arg at position 165 in human mitochondrial ME (position 151 in MEZ) is also critical for catalysis, and its modification disrupts malate binding without disrupting cofactor binding. While this Arg is highly conserved among eukaryotic species, it is less conserved among prokaryotes (**Figure 2B**). The substitution of Arg for Ala in this position for MEZ (Ala151) distinguishes it from the other structurally resolved large subunit prokaryotic *E. coli* ME, where the Arg is conserved (*E. coli* ME Arg157).

We solved the X-ray crystal structure of MEZ in its apo conformation. We attempted to soak the apo-MEZ crystals with high concentrations and to co-crystallize MEZ with malate, MgCl_2_ or MnCl_2,_ NAD(P)H and allosteric effectors. However, crystals showed no improvement in resolution and no observable electron density for any small molecules. To fully understand the structure of MEZ in the presence of its substrate, divalent metal and cofactor, we placed these components into the apo MEZ structure referencing the small molecule binding modes observed in the human mitochondrial ME structure (PDB: 1PJ2). We used Mn^2+^ as the divalent metal as MEZ has a preference for Mn^2+^ compared to Mg^2+^ [3]. After docking, we energy-minimized the structural complexes with Amber [30] before performing MD simulations. Because metals often present challenges in parameterization of classical molecular dynamics simulations, we initially simulated MEZ in the presence of malate and either NAD^*+*^ or NADP^*+*^ only. However, in the absence of Mn^2+^, we observed that malate binding is unstable in the active site (**Figure 4**) and dissociates within the first 10 ns of simulation.

**Figure 4.**
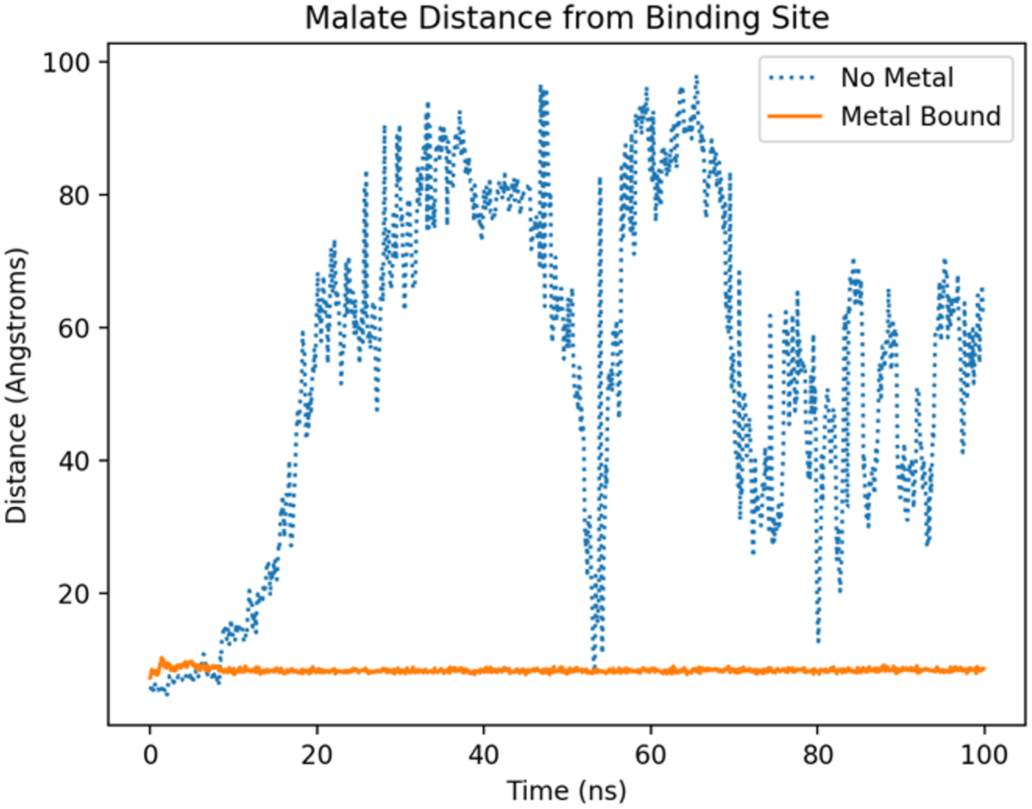
Divalent Cation Stabilizes Malate in Binding Site. In MD simulations of MEZ with malate and NAD^+^, the malate ligand exits the binding site within the first 10 ns of simulation and briefly returns and exits again around 55 ns. Comparatively, in simulations of MEZ with malate, NAD^+^, and the divalent metal Mn^2+^, the malate remains bound within the active site of MEZ for the duration of simulations.

After minimizing and equilibrating the structure of MEZ with Mn^2+^, malate and NAD(P)^+^ bound, we observe that within subunit A, the metal ion, malate and NAD(P)^+^ molecules, are not significantly displaced, and malate is stabilized by both the divalent metal and the nicotinamide group of NAD(P)^+^. However, within subunit B, we observed a greater degree of drift of the divalent metal and small molecules (**Figure 5**). In the presence of NAD^+^, subunit A retains contacts along the entire length of the dinucleotide, whereas the contacts of the adenine base with MEZ are lost within subunit B (**Figure 5A & Figure S4A**). The same is observed for NADP^+^, where the contacts of MEZ chain A with NADP^+^ form along the length of molecule, and in contrast, there is no contact between chain B and the adenine base, **Figure 5B & Figure S4B**. Even though the ribose 2’-phosphate forms no contacts with MEZ in chain B, NADP^+^ in chain B is only slightly displaced from NADP^+^ in chain A (**Figure 5**).

**Figure 5.**
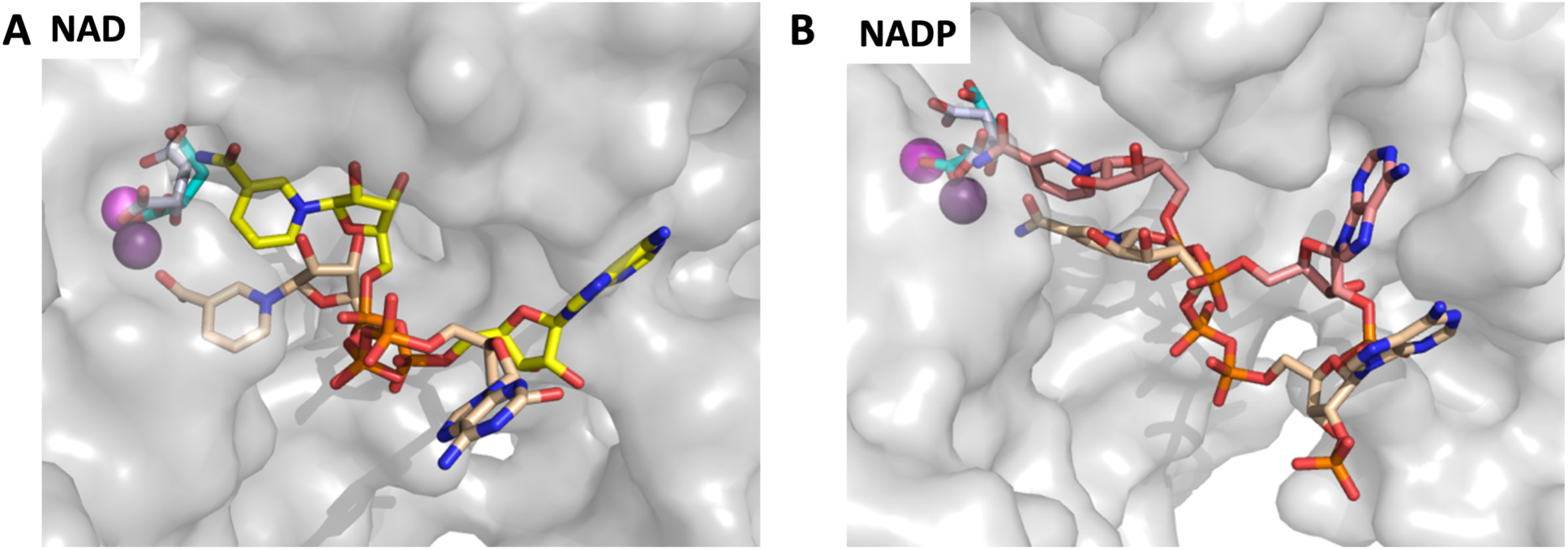
Different modes of NAD(P)^+^ binding after docking with Mn^2+^ and malate. Docked and minimized model showing superimposed subunits of MEZ (grey surface representation) the divalent metal (sphere) and small molecules (stick). **A**. After minimization, the left panel shows malate, Mn^2+^, NAD^+^, which are colored magenta, cyan and yellow, respectively, in chain A, and purple, grey and wheat in chain B. In chain A the molecules are in the correct orientation for catalysis, whereas in chain B, the molecules are no longer positioned to carry out the chemistry, as the nicotinamide group is not coordinated to malate, and in turn, malate to Mn^2+^. **B**. The right panel shows a similar result for NADP^+^ after minimization. In chain A, NADP^+^ (salmon) is positioned for catalysis but in chain B the NADP^+^ (wheat) nicotinamide group is oriented differently with respect to malate and Mn^2+^.

### Conformational Flexibility of MEZ

As discussed above, while some MEs use only NAD^*+*^ or NADP^*+*^ as an oxidizing agent to catalyze the conversion of malate to pyruvate, MEZ is among those MEs which can use either dinucleotide cofactor. To explore the NAD(P)^*+*^ binding modes, examine the potential cooperative dynamics between subunits within the dimer, and provide insight into the specific interactions that facilitate use of either NAD^*+*^ or NADP^*+*^ for catalysis, we ran MD simulations of MEZ with malate, NAD^*+*^ or NADP^*+*^, and Mn^2+^.

To evaluate and compare the overall flexibility of MEZ in its apo (open) and holo (closed) forms, we computed the root mean squared fluctuation (RMSF) of the αC atom for each residue and averaged the values over the five simulation repeats (**Figure S5A**). For domains A, B1 and B2 the RMSF values generally range between 1 and 3 Å and these regions are relatively stable compared to the rest of the protein chain. Domain D, and especially domain C, exhibit the greatest instability; residues in Domain D have RMSF values ranging from 1.5-3.5 Å while domain C has the broadest range from 1-7 Å for the apo MEZ simulation. When compared to RMSF values of other systems, domains C and D are particularly flexible; RMSF values computed from simulations of the highly compact and stable lysozyme are below 1 Å [31], while dimers of Amyloid β(1-40), a highly mobile and flexible protein range from 1-6 Å [33]. Interestingly, though binding might typically be expected to modestly stabilize fluctuations the addition of malate, Mn^2+^ and NAD(P)^*+*^ to the simulations had no impact on the overall stability of the MEZ protein backbone, apart from some reduction in computed αC RMSF values for residues between Gln355 and Ala430 (**Figure S5B**). This is consistent with our empirical observations that soaking or co-crystallizing substrates, cofactors and metals with MEZ failed to improve the diffraction quality of our crystals.

### Fluctuations Between Open and Closed States

Previous studies of MEZ homologs suggest that MEs can adopt open and closed states [13, 31]. In our simulations we also observe expansion and contraction of the MEZ active site, which can be tracked by recording the distance between αC atoms on two neighboring helices (**Figure S6**). Here, we evaluated the relative stability of cofactor binding in neighboring subunits and whether there is an interdependence of open and closed states within the dimer.

Among the five simulations of MEZ in the presence of Mn^2+^, malate and NADP^+^, the subunits adopt both closed and open states (all 5 simulations are shown in **Figure S6**), with subunits in some simulations remaining stably in one state or the other, but in other cases, multiple transitions occur. In simulations where the cofactor remains stably bound in an active orientation with the nicotinamide group near the malate and Mn^2+^ (Simulations 1, 3, 5), the subunits stably adopt opposite forms or in the case of Simulation 5 appear to rapidly fluctuate between the open and closed states. In the case of MEZ with Mn^2+^, malate and NAD^+^, the subunits also transition between open and closed states. Further, for simulations where the NAD^+^ cofactor is stably bound to both subunits in a catalytically relevant orientation (Simulations 3, 5), the subunits within the dimer adopt opposite open-closed forms.

Because typically molecular mechanics simulations cannot simulate catalytic mechanisms, specifically the making and breaking of bonds and electron transfer, these simulations do not provide direct insight into the mechanism of malate turnover. However, we can draw general conclusions regarding the global dynamics of the protein in association with its substrate and cofactors. All simulations showed a fluctuation between the open and closed forms. Furthermore, in simulations with NAD(P)^*+*^, malate and Mn^2+^, where active sites from both subunits are occupied by cofactor, substrate and divalent cation, the data suggests that within the dimer, the closed ‘active’ form is only stabilized when the neighboring subunit stably adopts an open conformation, even though the two active sites within the dimer are not in close proximity to one another (**Figure S7**). These results are further supported by the kinetic data above, which suggests that MEZ is allosterically regulated by the substrate malate.

### NAD(P)^+^ Co-factor Binding Modes

To identify the residues that participate in cofactor binding and that specifically facilitate the binding of both NAD^+^ and NADP^+^, we selected the simulations for which the cofactor binding was most stable for both MEZ subunits and used the final simulation frame as the reference for analysis. To compare relative stability amongst simulations, we computed the average RMSF for each cofactor atom in both subunits. For MEZ with Mn^2+^, malate and NAD^+^, we selected Simulation 3; while for MEZ with Mn^2+^, malate and NADP^+^, we selected Simulation 1 (**Figure S6**).

With Mn^2+^, malate and NAD^+^, we observe two stable conformations of MEZ: one elongated and the other collapsed (**Figure 6B**). Indeed, multiple conformations of NAD(H) have been observed and described previously in crystallographic, solution NMR, and MD studies [31-34]. Specifically, the collapsed form is commonly observed for unbound NAD(H) in solution [35], but has also been observed in holo structures, including in the exo-site of human mitochondrial ME (PDB ID:1PJ3, 1PJL [36, 37]). To identify additional examples of enzymes in complex with the collapsed form of NAD^+^, we retrieved all instances of NAD^+^ molecules deposited in the PDB (a total of 4,163 sets of coordinates) and computed Tanimoto scores with OpenEye Scientific’s Shape Toolkit using the collapsed form we observe in our MEZ simulations as the reference structure. The three highest scoring NAD^+^ molecules (with the most similar conformation to the collapsed NAD^+^ in MEZ) were observed in *Apylsia California* ADP ribosyl cyclase, which cyclizes NAD^+^ into cyclic-ADP-ribose (PDB ID: 3ZWM [38]), as well as prokaryotic CBS domain proteins (PDB ID: 2RC3, 4FRY), whereby the domain regulates enzyme and transport activities in response to adenosyl group binding.

**Figure 6.**
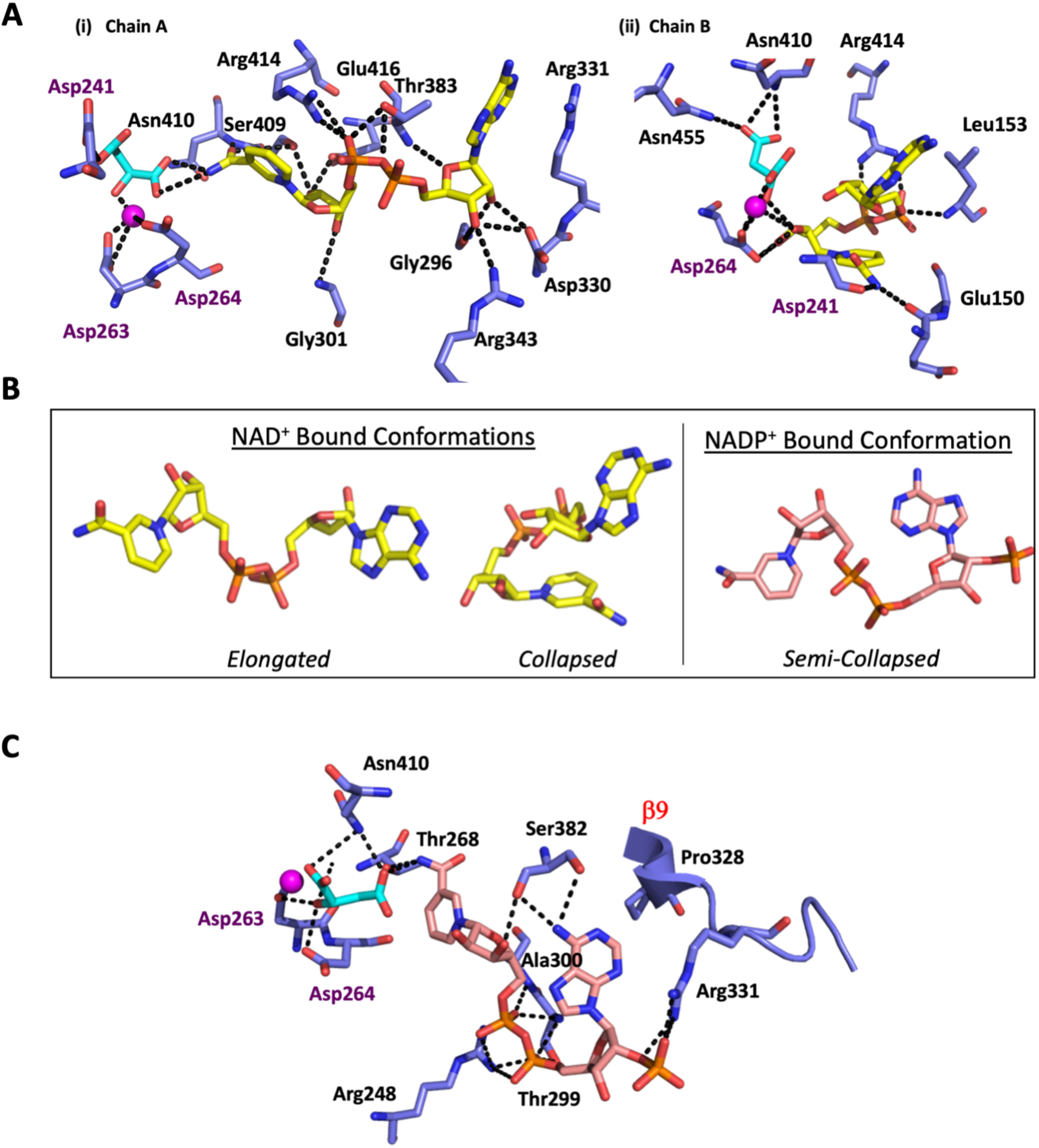
Putative NAD(P)^+^ Binding Modes in MEZ after MD simulations. **A**. Putative NAD^+^ binding modes in MEZ. In simulations of MEZ (blue) with NAD^+^ (yellow), Mn^2+^ (magenta), and malate (cyan), the NAD^+^ cofactor adopts two distinct binding modes. In **i**) the NAD^+^ extends along the pocket in an elongated conformation. The nicotinamide interacts with the malate while making polar contacts with many MEZ residues. In **ii**) the NAD^+^ cofactor is folded over in a collapsed conformation and occupies a much more limited space of the MEZ binding pocket. The nicotinamide interacts with MEZ residues, Mn^2+^ and malate substrate, and thus this likely represents an inactive bound conformation. **B**. Bound cofactor conformations. Representative conformations of the NAD^+^ (yellow) and NADP^+^ (pink) binding modes in simulations of MEZ with Mn^2+^ and malate. The conformations were selected by taking the final frame of the simulation trajectories where the cofactors were most stably bound. The elongated form of NAD^+^ represents an active conformation while the collapsed form of NAD^+^ occupies the binding site in an inactive formation. For NADP^+^ simulations, the cofactor occupies a single semi-collapsed, active form with the nicotinamide oriented near the malate substrate. **C**. Putative NADP^+^ binding mode in MEZ. In simulations of MEZ with NADP^+^ (pink), Mn^2+^, and malate, NADP^+^ makes several polar interactions with residues in the MEZ binding pocket and with malate. The distinct kinking of the β9 strand in MEZ by Pro328 may favorably position Arg331 for interaction with the NADP^+^ 2’-phosphate group.

Here we suggest that the collapsed form of NAD^+^ in complex with MEZ, represents an inactive, albeit stable conformation; the nicotinamide group is oriented away from the malate substrate and Mn^2+^ cation, and instead makes polar contacts with the backbone of Glu150 (**Figure 6A (ii)**). This form is also stabilized by polar contacts with malate, Asp264, Arg414, and the backbone of Leu153 and Asp241. As noted previously, human mitochondrial ME (PDB: 1PJ3) as well as all other structurally resolved large subunit MEs, have a conserved ‘GER (Glu163-Arg165)’ motif in the active site, corresponding to residues 149-151 in MEZ. In MEZ (and other small subunit prokaryotic MEs), this motif is substituted for residues ‘AEA,’ beginning with Ala151. Given that the collapsed form of NAD^+^ is proximal to Ala151, it is possible that the presence of an Arg residue at this position could interrupt or perturb the stability of the collapsed conformation in the MEZ active site. We speculate that this collapsed form of NAD^+^ may be unique among non-’GER’ containing large subunit MEs. The elongated conformation of NAD^+^ appears to adopt a catalytically relevant orientation (**Figure 6A (i)**), with the nicotinamide group oriented toward the malate and Mn^2+^ cation. This active form is stabilized by several polar interactions with the malate substrate and sidechains of Arg343, Asp330, Thr383, Ser409, Arg414 as well as the backbone of Gly301 and Asn410 (**Figure S2B**), and an additional interaction of Arg331 sidechain with the aromatic ring of the adenine base.

By comparison, in simulations of MEZ with Mn^2+^, malate and NADP^+^, the cofactor adopts a semi-collapsed form (**Figure 6B**) which appears to be well oriented for catalysis with the nicotinamide group located near the malate substrate and Mn^2+^ cation (**Figure 6C**). NADP^+^ makes several polar contacts with malate as well as sidechain and backbone atoms of the surrounding residues. The ligand is stabilized by interactions with Arg248, Thr268, Thr299, Ala300, Arg331, and Ser382. Notably the kinking of β9 by Pro328 (**Figure S3**), helps orient Arg331 such that it can stabilize the additional 2’-phosphate group on the ribose (absent in NAD^+^). Tanimoto analysis was performed using the semi-collapsed structure of NADP^+^ and all structures of NADP^+^ in the PDB (3164 sets of coordinates). Interestingly, the three top-scoring NADP^+^ molecules are found in structures of Mtb nicotinamide-mononucleotide adenylyltransferase (NadD) (PDB ID: 4S1O) and *Mycobacterium abscessus* NadD (PDB ID: 4YMI & 5VIR). These results suggest that the semi-collapsed NADP^+^ form observed in MEZ MD simulations is biologically relevant.

## Discussion

Many bacterial species have multiple copies of genes encoding for MEs [15, 19, 39], which underscores this enzyme’s physiological importance in central carbon metabolism as well as in the provision of NAD(P)H for anabolic processes such as fatty acid biosynthesis. Several eukaryotes also possess multiple ME isoforms, which are often distinct from each other in their cofactor preference. For example, humans possess three MEs; NADP-dependent ME is found in the cytosol while the other two, are localized to the mitochondria [14]. Analogously, photosynthetic eukaryotes possess both plastidic C4-associated tetrameric ME with a role in photosynthesis as well as a non-C4 dimeric ME isoform [12, 40, 41]. By contrast, Mtb and other are mycobacteria are reliant on only one ME and as such, will require flexible mechanisms of regulation in order to satisfy metabolic requirements.

### Unlike other characterized large subunit MEs, MEZ forms a stable dimer

Generally, eukaryotic MEs are observed as either a stable tetramer or in a dimer-tetramer equilibrium, wherein substrate binding or allosteric activators drive the formation of the higher-order tetrameric state [12, 40, 42, 43]. Structurally, stabilization of the tetramer for larger subunit MEs is facilitated by the presence of a C-or N-terminal extension [12, 14]. Mutations in the C-terminal extensions of pigeon ME and human cytosolic ME disrupt formation of the tetramer assembly [16, 17]. In photosynthetic eukaryotes, the N-terminal extension is long (50-60 residues) when compared to the moderately extended C-termini of non-photosynthetic eukaryotes (20-30 residues). Structurally, both the N- and C-termini localize at the dimer and tetramer interfaces (**Figure 1B**). In non-photosynthetic eukaryotes, the C-terminus extends up and away from the dimer along the middle of the solvent facing region of the adjacent dimer subunit, while in photosynthetic eukaryotes, the N-terminus forms a β-strand that interacts with the neighboring dimers’ N-terminus to form a pair of antiparallel β-sheets at the vertex of the tetramer interface (**Figure 1B**). *E. coli* ME, MEZ’s closest structural homolog, has modest 17-residue and 7-residue extensions at the N- and C-termini, respectively (**Figure S2B**). Notably in *E. coli*, these N- and C-terminal extensions form β-strands, which come together to form a parallel β-sheet near the canonical dimer-dimer interface (**Figure 1B**). Both prokaryotic large subunit MEs, *E. coli* ME and MEZ, crystallized as a dimer; however SEC of *E. coli* ME suggests it assembles as a tetramer [19]. In summary, MEZ has neither an N-nor C-terminal extension, the canonical ME tetramer is not observed in the X-ray crystal structure, and in solution we confirmed that MEZ is a stable dimer (**Figure 3A**).

### MEZ may be partially regulated by pH

The regulation of MEs by pH has been studied in other organisms. In eukaryotic species, it has been observed that acidic pH drives the dissociation of the tetramer/dimer equilibrium toward dimeric and monomeric species. Disruption of tetramer formation has been linked to substrate inhibition of ME by malate at a pH of less than 7.0 [25, 26, 44-46]. Photosynthetic C4-ME is inhibited by malate in a pH dependent manner when the stromal pH drops to 7.0-7.5 from pH 8.0 [12]. The molecular basis of this inhibition was shown to be related to destabilization of the higher order oligomeric state [12]. In our DSF studies, we observe that MEZ is stabilized at lower pH; however, by SEC the oligomeric state persists as a dimer at both pH 7.5 (**Figure 3A**) and pH 6.0 (data not shown). This may have biological relevance as *mez* is part of the *phoPR* regulon, which is upregulated at low pH [8]. In support of this, we show biochemically that the catalytic efficiency of MEZ improves at lower pH.

### MEZ is allosterically regulated by fumarate and succinate

Our biochemical analysis of MEZ suggests that like many eukaryotic MEs, MEZ operates with cooperative kinetics with respect to malate [21, 22]. In the case of human mitochondrial and *Ascaris suum* MEs, fumarate acts as an allosteric activator and supplants the cooperative kinetic effects of malate binding [25, 27]. For human mitochondrial ME, the fumarate binds at two symmetric allosteric sites at the dimer interface; these two sites are coordinated by Arg67 and Arg91 residues from each subunit. While our MEZ structure has no ligands bound at these sites, the structurally corresponding residues, Arg51 and Arg75 are organized in a similar configuration at the dimer interface, suggesting that binding of ligands at the two symmetric allosteric sites may also disrupt cooperative binding of malate in MEZ. Further, our kinetic data shows that decarboxylating activity is enhanced in the presence of fumarate, succinate and alanine and inhibited by oxaloacetate (**Figure 3B**). Finally, MEZ activity is enhanced by fumarate more than succinate, which likely stems from higher energetic favorability in binding the more rigid fumarate due to reduced entropic costs or from the fact that fumarate is more easily deionized at low pH in comparison to succinate.

Most of the characterized MEs are allosterically controlled. Indeed studies of the homologous *E. coli* ME demonstrated allosteric upregulation by aspartate and downregulation by oxaloacetate, CoA, palmitoyl-CoA, and acetyl-phosphate [19]. However, fumarate, glutamate, succinate, glycine and alanine had no effect on enzyme activity. Metabolically it is advantageous that fumarate and succinate, both products of the TCA cycle immediately upstream of malate, regulate MEZ (**Figure S1**). Similarly, it follows that oxaloacetate, the metabolite downstream of malate, is inhibitory. The observed modulation of enzyme activity by alanine could potentially be regulated through the activity of alanine dehydrogenase (ALD), which converts pyruvate (the product of MEZ) to alanine along with the concomitant oxidation of NADH to NAD^+^. Like MEZ, the gene for ALD is upregulated under hypoxic conditions [47], leading one to postulate that MEZ may play a role in preserving redox balance and supplying NAD(P)H when ALD activity increases.

### The plasticity of MEZ: its biological relevance and druggability

Generally speaking, it has been observed that structural protein disorder is associated with either 1) increased promiscuity and/or 2) a disorder-to-order transition upon protein-ligand interactions, or 3) a disorder-to-order transition upon protein-protein interactions [39]. Given that we do not observe a decrease in disorder in our simulations nor an increase in MEZ thermal stability in the presence of cofactors and substrate (**Figure 3C**), there is little evidence to support a disorder-to-order transition upon ligand binding. We do not know whether MEZ forms protein-protein interactions that facilitate a transition to a more ordered state. However, a recent study demonstrated that cytosolic human ME1 activates and forms a stable hetero-complex with 6-phosphogluconate dehydrogenase (6GPD), which notably also generates NADPH to potentially maintain redox homeostasis [48]. Mtb has an analogous coenzyme F420-dependent 6-phosphogluconate dehydrogenase (FGD1), which generates reduced coenzyme F420. While FGD1 is structurally distinct from human 6GPD [49], MEZ may interact with FGD1 or have other protein-interaction partners. Ultimately, we posit that the structural plasticity of MEZ facilitates cofactor or substrate promiscuity and allosteric regulation. These features allow MEZ to generate reductants for lipid biosynthesis and the maintenance of redox homeostasis rapidly in response to metabolic and environmental stressors, which Mtb experiences in the human host.

As MEZ plays a role in lipid biosynthesis [3], this enzyme presents a potential anti-TB drug target. Similarly, human MEs have also been identified as potential targets for developing anti-cancer therapeutics [24, 50, 51]. Importantly, there are distinct structural differences between MEZ and the human ME homologs. In our simulations, we observe that the NAD^+^ cofactor can adopt multiple stable binding modes (**Figure 6**). For the compact form of MEZ, the ribose group near the NAD^+^ adenine motif forms polar contacts with the Mn^2+^ ion and malate substrate, but the nicotinamide group is oriented away from the active site and thus, is not suitably positioned for catalysis (**Figure 6A (ii)**). In order to accommodate this orientation, MEZ adopts a semi-open conformation wherein the β-hairpin turn (β3/β4) of residues 150-154 is pushed further away from the binding pocket as its backbone makes interactions with the compact NAD^+^ molecule. Notably, in other ME structures such as human mitochondrial ME, the residues at the apex of this β-hairpin form a helix that is flanked by the Arg165 residue, which anchors this turn to the ME active site by forming polar contacts with malate (PDB ID: 1PJ2). In MEZ, Arg165 is substituted for an alanine (Ala151), and as the β-hairpin no longer possesses a positively charged sidechain to coordinate the malate substrate, the β-hairpin has greater flexibility. Indeed, this flexibility is observed in the apo MEZ structure as the residues that form the β-hairpin are random coil in Chain A, but an ordered β-hairpin in chains B, C & D. We suspect that the stable binding of NAD^+^ in a compact form is unique to MEZ and other prokaryotic MEs that lack the conserved Arg165 residue. Interestingly, this compact NAD^+^ conformation is similar to cyclic-ADP-ribose [38]. As MEZ can accommodate the compact form due to the structural differences compared to human MEs, an inactive cyclized NAD derivative may be an excellent starting point to design inhibitors of MEZ, which would not target human MEs. The role of MEZ in the Mtb life cycle and its unique structural qualities presents an appealing opportunity for the design of selective inhibitors, and warrants further exploration.

## Methods

### MEZ purification

MEZ was purified as described [3]. Briefly a pNIC28-BSA4 plasmid containing the MEZ gene with an N-terminal His-tag was transformed into BL21(DE3) cells and grown in LB media containing 50 μg/ml kanamycin at 37°C. Once the cells reached 0.6-0.8 OD, expression was induced with 1mM IPTG and the temperature was reduced to 18 °C for approximately 18 hours. Cells were harvested by centrifugation for 30 minutes at 5000 rpm. Pellets were washed with STE buffer (10 mM Tris-HCl, pH8, 100 mM NaCl, 1 mM EDTA) and stored at −20°C. Cell pellets were resuspended in Buffer A (50 mM Tris pH 7.6, 500 mM NaCl, 50 mM Imidazole, 0.14 mM BME (β-mercaptoethanol), 5mM MgCl_2_, 4% glycerol) containing lysozyme and 1 mM PMSF (phenylmethylsulfonyl fluoride). Following sonication and centrifugation to remove debris, the cell lysate was loaded onto a Ni^2+^-bound His-Trap column and washed with several column volumes of Buffer A, followed by a high salt wash (three column volumes of Buffer A with 2M NaCl). The MEZ was eluted with a gradient of Buffer A and Buffer A containing 500mM Imidazole. MEZ containing fractions were identified via SDS-PAGE analysis and collected and pooled. For DSF and size exclusion chromatography, MEZ was dialyzed into Buffer A overnight at 4°C.

### MEZ crystallization

Prior to crystallization studies, MEZ was concentrated and dialyzed (Amicon, 30 kDa MWCO) into 50mM trisodium Citrate-citric acid, 100mM NaCl, pH 6.0, per the results of a pre-crystallization DSF buffer screen performed using the Meltdown program [52], wherein MEZ exhibited a trend of greater thermal stability at lower pH (pHs 5.5-10 were tested). Notably, prior attempts at crystallization (before identification of an optimal protein buffer and pH), yielded few diffracting crystals and none were reproducible. Crystallization conditions were identified via a collection of Hampton Research sparse matrix crystal screens. Crystal hits were optimized to produce diffraction quality crystals. Notably, protein concentration and drop sizes were all varied. Cryo-conditions crystal seeding, in-situ proteolysis, dehydration, additive screens and the addition of small organic compounds including alanine, glycine, tartronic acid, malate and cofactors NADH and NADPH (2-10mM) via soaking and co-crystallization were all explored. Of the more than 2000 crystals that were harvested, most diffracted no better than 8.0 Å and just two crystals diffracted to 4.0 Å or better.

The crystal condition that gave the highest resolution diffracting crystal was 0.2M NaCl, 0.1M HEPES, pH 7.5, 2mM alanine, 12% PEG 8000 and the crystal was cryo-frozen in 20% glycerol. The diffraction data set was collected at 70K to a resolution of 3.6 Å at ALS and was indexed, integrated and scaled in MOSFLM [53]. A solution was generated via molecular replacement using Phaser [54], utilizing a dimer search model generated by iTasser [55] from the structure of human ME (PDB ID: 2AW5, chains B&C). The model was refined in Coot and phenix.refine [54, 56].

### MEZ Enzyme Assays

MEZ enzyme activity was assayed as described [3]. Briefly, MEZ enzyme activity was measured at 30 °C by spectrophotometrically measuring the NAD(P)H formation at 340 nm. The reaction mixture contained: 100 mM Tris, 100 mM malate, 0.5 mM MnCl_2_, 1.6 mM NAD(P)^+^ at pH 7.4 or pH 6.6. The kinetic constants were determined by varying the concentration of malate (0.25 - 100 mM) or NAD(P)^+^ (0.05-1.6 mM) while leaving other components at saturation concentrations. When different compounds were tested as potential inhibitors or activators of enzyme activity, MEZ forward activities (using NAD^+^ as a cofactor) were measured at pH 7.4 in the absence or presence of 0.5, 1.0, or 2.0 mM of each effector (oxaloacetate, succinate, fumarate, pyruvate, aspartate, glutamate, acetate or alanine) whilst the malate concentration was maintained at 4 mM, approximately one-fifth of the *K*_*0*.*5*_, malate value. The results are presented as the percentages of activity in the presence of the effectors in relation to the activity measured in the absence of the metabolites (100%).

### Protein Thermal Stability Measurements

Protein thermal stability was determined using differential scanning fluorimetry (DSF). For DSF measurements, proteins were incubated with 25 or 40 μM SYPRO orange dye in 20 mM sodium phosphate (pH 7.4), 150 mM NaCl. Samples were heated from 25 °C to 96 °C at 1 °C/min using an Mx3005P qPCR machine (Agilent Technologies). The dye was excited at 492 nm, and fluorescence emission was monitored at 610 nm. Melting curves were obtained in duplicate, and each experiment was conducted independently three times. Melting temperatures (*T*_*m*_s) were determined using nonlinear regression to determine melting-curve inflection points.

### Size exclusion chromatography

Size exclusion chromatography for MEZ was run on a S200 HiLoad 16/600 column at 0.5mL/min with protein standards from Bio-Rad to determine the oligomeric state of MEZ in solution. MEZ was run in its apo form, or with the addition of 100 molar excess of fumarate, succinate and alanine, with 2mM of the small molecule in the running buffer.

### Molecular Dynamics Simulations

To prepare MEZ for simulations, a dimer of the most complete chain B was generated by aligning a duplicate copy of chain B with its partner, chain A, in PyMol (DeLano Scientific, CA) and submitting the B/B dimer to pdbfixer [57] to model in missing atoms, residues and loops using default protonation states at pH 7.0. Malate, NAD^+^ and NADP^+^ were manually docked in PyMol so as to mimic binding modes observed in MEZ’s closest structural homolog by sequence, human mitochondrial ME (PDB ID: 1PJ2).

Cofactors NAD^+^ and NADP^+^ were prepared using previously optimized Amber parameters [58, 59]. The Mn^2+^ metal site parameters were built using Amber’s MCPB.py program [60]. Malate and Mn^2+^ were prepared using antechamber from AmberTools18 [30] with GAFF version 1.7 and AM1-BCC charges. In the MCPB.py input file, the terminal oxygens of residues Asp241 and Asp264 were listed as having bonded pairs with the Mn^2+^ ion and the malate ligand was identified as a non-amino acid included in the metal complex. Geometry optimization and force constant calculations were performed with input files generated by MCPB.py using Gaussian09. Output parameters were evaluated with parmed [61] and satisfied the criteria outlined in the MCPB.py tutorial.

Following this, each system was solvated using tleap from AmberTools18 [30] with a 10 Å rectangular box of TIP3P water, extending from the surface of the protein to the box edge with sodium ions added to neutralize the charge of the system. Minimization proceeded using sander from Amber14 with steepest descents running for 20000 steps, followed by heating from 100 to 300 K with a constant volume for 25000 time steps of 2 fs. Equilibration continued using sander for 500000 time steps of 2 fs under constant pressure with positional restraints initially applied on all non-water atoms at 50 kcal/mol/Å^2^ and progressively increased in increments of 5 kcal/mol/Å^2^ over ten 50000-step segments. The resulting topology and coordinate files for each system were used as inputs for further MD simulations. Production simulations were executed in OpenMM 7.1.0 [62] using a Langevin integrator with a 2 fs time step and a friction coefficient of 10 ps–1. For each of four systems (MEZ only, MEZ/malate/NAD^+^, MEZ/Mn^2+^/malate/NAD^+^, and MEZ/Mn^2+^/malate/NADP^+^), five 100 ns simulations were initiated with random starting velocities.

### Molecular Dynamics Analysis

Analysis of trajectory data was performed using MDTraj 1.9.3. In order to systematically track the stability of malate in the binding site, we computed the distance between C1 atom of malate and the αC atom of Asn455 for the duration of each simulation. Similarly, the relative positioning of NAD(P)^+^ in relation to the active site was tracked by recording the distance between the C4 atom of the nicotinamide group of NAD(P)^+^, which is reduced to form NAD(P)H, and the αC atom of Asn455. To quantify differences in the conformational stability of MEZ alone or in complex with malate and NAD(P)^+^, we computed and averaged the RMSF values of each αC atom in the protein backbone for all simulations and plotted them as shown in (**Supplemental Figure 5**). Visual inspection of the trajectories revealed open and closed states for the MEZ subunits; in order to assign the relative state, we tracked the relative proximity of two neighboring helices by computing the distance between αC atoms of Pro244 and Ala300 (**Supplemental Figure 6**). For analysis of NAD(P)^+^ binding modes, we identified one simulation among the five replicates of MEZ/Mn^2+^/malate/NAD^+^ and MEZ/Mn^2+^/malate/NADP^+^ for which the cofactor was most stable. Sustained binding of the cofactor in the active site of MEZ was assessed by computing the distance between the NAD(P)^+^ carbon atom at position 4 of the nicotinamide group and the αC atom of a stable residue in the binding pocket, Asn455. Relative stability of the cofactor amongst the replicates was imputed by comparing the RMSF for each atom in the NAD^+^ or NADP^+^ ligand; the final frame was used to examine the contacts between the protein and each cofactor.

## Accession Numbers

Mtb ME (MEZ): P9WK25 PDB ID code: 6URF.

## Abbreviations

6GPD: 6-phosphogluconate dehydrogenase
ADP: adenosine diphosphate
ALD: alanine dehydrogenase
BME: β-mercaptoethanol
DSF: differential scanning fluorimetry
FGD1: F420-dependent 6-phosphogluconate dehydrogenase
HIV: human immunodeficiency virus
IPTG: isopropyl β-d-1-thiogalactopyranoside kDa – kilodalton
MD: molecular dynamics ME – malic enzyme
MEZ: *Mycobacterium tuberculosis* malic enzyme
Mtb: *Mycobacterium tuberculosis*
NAD(P)^+^ / NAD(P)H: nicotinamide adenine dinucleotide (phosphate)
NadD: mononucleotide adenylyltransferase
NMR: nuclear magnetic resonance
PMSF: phenylmethylsulfonyl fluoride
PDB: protein data bank
rmsd: root mean square deviation
RMSF: root mean square fluctuation
SEC: size exclusion chromatography
TAG: triacylglycerol
TB: Tuberculosis
TCA: tricarboxylic acid cycle

## Acknowledgements

We would like to thank Tom Mendum for useful discussions and Tom Poulos and Jose Artiga for critical reading of the manuscript. We thank the Advanced Light Source at Berkeley National Laboratories (ALS) and the Stanford Synchrotron Radiation Lightsource (SSRL) for their invaluable help in data collection, and The Triton Shared Computing Cluster (TCSS) at the San Diego Supercomputing Center (SSDC) for their computing support.

